# Comprehensive analysis of tumour initiation, spatial and temporal progression under multiple lines of treatment

**DOI:** 10.1101/508127

**Authors:** Ignaty Leshchiner, Dimitri Livitz, Justin F. Gainor, Daniel Rosebrock, Oliver Spiro, Aina Martinez, Edmund Mroz, Jessica J. Lin, Chip Stewart, Jaegil Kim, Liudmila Elagina, Ivana Bozic, Mari Mino-Kenudson, Marguerite Rooney, Sai-Hong Ignatius Ou, Catherine J. Wu, James W. Rocco, Jeffrey A. Engelman, Alice T. Shaw, Gad Getz

## Abstract

Driver mutations alter cells from normal to cancer through several evolutionary epochs: premalignancy, early malignancy, subclonal diversification, metastasis and resistance to therapy. Later stages of disease can be explored through analyzing multiple samples collected longitudinally, on or between successive treatments, and finally at time of autopsy. It is also possible to study earlier stages of cancer development through probabilistic reconstruction of developmental trajectories based on mutational information preserved in the genome. Here we present a suite of tools, called Phylogic N-Dimensional with Timing (PhylogicNDT), that statistically model phylogenetic and evolutionary trajectories based on mutation and copy-number data representing samples taken at single or multiple time points. PhylogicNDT can be used to infer: (i) the order of clonal driver events (including in pre-cancerous stages); (ii) subclonal populations of cells and their phylogenetic relationships; and (iii) cell population dynamics. We demonstrate the use of PhylogicNDT by applying it to whole-exome and whole-genome data of 498 lung adenocarcinoma samples (434 previously available and 64 of newly generated data). We identify significantly different progression trajectories across subtypes of lung adenocarcinoma (*EGFR* mutant, *KRAS* mutant, fusion-driven and *EGFR/KRAS* wild type cancers). In addition, we study the progression of fusion-driven lung cancer in 21 patients by analyzing samples from multiple timepoints during treatment with 1st and next generation tyrosine kinase inhibitors. We characterize their subclonal diversification, dynamics, selection, and changes in mutational signatures and neoantigen load. This methodology will enable a systematic study of tumour initiation, progression and resistance across cancer types and therapies.

## Introduction

Studying tumour heterogeneity, clonal structure and dynamics is becoming an increasingly important clinical and research topic. Many new and current cancer therapies show efficacy at the start of treatment but stop working rapidly, leading to disease progression ^1^. Understanding the processes of acquired and intrinsic resistance to therapy, evasion of the host immune system, and metastatic spread is important to advance clinical care.

It is widely understood that tumours arise through successive accumulation of somatic aberrations in the genome of individual cells. These aberrations can sometimes confer growth advantage and result in clonal expansions, further progressing towards malignancy. The overall order and accumulation of acquired events can be described as a developmental trajectory with some clonal subpopulations being selected out and thus becoming extinct in the process. During tumour progression with and without treatment, cancer cells often undergo successive population bottlenecks and evolve to maintain the fittest clones under the current conditions. Multiple sites within a tumour will often have distinct subclonal compositions, depending on both genetic and micro-environmental factors ^2,3^. These conditions lead to widespread spatial clonal heterogeneity within and between lesions. Over time, these different cancer cell populations can become substantially distinct in their genetics and biological behavior even though they are still all genetically related.

In several tissue types, there are well-known premalignant or clonally expanded lesions that do not necessarily progress to cancer (e.g. Barrett’s esophagus, colorectal polyps etc.). The number of people that exhibit such detectable premalignant expansions by far exceed the number of patients that eventually progressed to cancer ^4,5^. It is possible that that there is a difference in the trajectories of these lesions and they might contain events that hinder progression to malignancy (i.e., “dead-end” trajectories).

If we were able to follow this progression route with some level of certainty and understand how these trajectories control the fate of the developing clone, this could lead to improved methods for outcome prediction, new early detection and cancer prevention approaches, as well as inform treatment decisions. Recent methodological developments allow exploration of these types of questions using sequencing data to a level of detail not available previously^6,7–11,12,13^. Both our and others work^6,7,5,11,14,15,16,17,18,19,20,21^ explored individual aspects of subclonal diversification and tumour progression in several different tumour types ^5–7,9,22,23^ and developed methods to computationally reconstruct absolute copy number^24,25^, subclonal structure of cell populations^5,26,21^ and population structure changes between samples^27,28,29,5,26,6^.

Several approaches also explored the possibility to time (i.e. order) early events, including ones that occurred prior to malignant transformation, by using evidence obtained from the amplification of mutated alleles (i.e. mutational multiplicity) ^30,31^. Several studies^13,30,31,32^, proposed a method to estimate the relative order of gains and copy-neutral loss-of-heterozygosity events by comparing the rates of multiplicity 1 and 2 mutations in whole-genome sequencing data. Other groups instead developed aggregate progression models based on assumptions of causality and event co-occurrence ^33,34^. More systematic experimental efforts were accomplished in some tumour types by studying data from premalignant lesions and manually compiling these progression models ^5,35,36^. Several recent studies explored intratumour heterogeneity in primary and later stage tumours, including in lung and kidney cancer (TRACERx)^23,37,38,39^.

It is important to appreciate that methods that attempt to reconstruct tumour evolutionary history based on sequencing data have only a limited view into the actual life history of the tumour. Moreover, measurements of copy number, mutation coverage and allele frequency and any other redout from next generation sequencing data inherently contain uncertainties that need to be properly accounted for and propagated to downstream analyses. These uncertainties, including random and systematic biases as well as limited discovery power, affect the modeling of all aspects of tumour development, including trajectories, phylogenetic relationships, clonal dynamics, growth rates, gene expression and mutational processes. Therefore, all estimations need to be performed in a consistent probabilistic manner. As we show below, obtaining multiple samples from a patient at different timepoints during tumour progression greatly increases the amount of information and resolution of such analyses.

Here, we report and apply a novel comprehensive package of tightly integrated tools, *PhylogicNDT*, for subclonal analysis, subclonal dynamics, event ordering, and timing, with primary focus of jointly modeling behavior of many samples from the same patient (both WES and/or WGS data). *PhylogicNDT*, extends our previous approaches ^5,7,9,19,26^ that were limited to analyzing of a single or a pair of samples per patient and enables joint and broader analysis of dozens of samples from a single patient, including WES and WGS data, generated from tissue samples, blood biopsies, autopsy samples, and cell lines. The package introduces significant algorithmic and methodological improvements, including novel tools (e.g. for tree building, clonal kinetics analysis and timing of somatic events in individual tumours and patient cohorts) that carefully and explicitly account for uncertainties in sequencing data and propagating posterior probabilities through the various analysis steps.

We apply the *PhylogicNDT* package to a cohort of lung adenocarcinomas, including multiple tumour samples per patient (up to 13) from 21 patients. Lung adenocarcinoma (LUAD) provides a particularly useful clinical scenario to investigate tumour evolution over time, including subclonal diversification, dynamics, and selection as well as changes in mutational signatures and neoantigen load associated with targeted therapies (1st and next, 2nd and 3rd, generation TKI treatment). LUADs driven by *ALK* fusion events are initially highly susceptible to treatment with TKIs, but inevitably recur with outgrowth of treatment-resistant cancer cells^40,41^. The recognition of this kinase activity offered a new opportunity for drug development and treatment, as these cells are highly susceptible and sensitive to tyrosine kinase inhibitors (TKIs) including crizotinib, ceritinib, alectinib, brigatinib and lorlatinib ^42^. The inevitable outgrowth of resistant tumour cells is most often due to secondary re-activating mutations in the ALK kinase domain as well as fusion gene amplification or other signaling pathways activation ^22,43,44^. Although analysis of post-progression biopsy specimens has proven valuable in facilitating a greater understanding of molecular mechanisms of crizotinib resistance ^22^, better characterization of the identity, order, and timing of the driver mutations required for acquired resistance is needed in order to develop more effective and long-lasting therapies.

Additionally, we applied PhylogicNDT to study different developmental trajectories (at both the individual patient level and cohort level) in the fusion-driven subtype of lung adenocarcinoma and other known clinical subtypes, such as KRAS- and EGFR-mutated tumours.

More generally, understanding and mapping the path (trajectory) which a normal cell takes to reach the malignant state and advance to resistance could lead to clinical and biological findings, explain differences between cancer subtypes, help predict treatment response and progression, develop faithful animal cancer models, and devise early detection techniques. This methodology can be applied to study tumour initiation, progression and resistance across cancer.

## Results

### Comprehensive analysis of tumour progression on single patient and cohort level

The PhylogicNDT package provides tools for integrated analysis of tumor progression from next-generation sequencing (NGS) data and contains the following components (Figure 1, Supp. Figure 1): (i) *Clustering* - used to identify clusters of mutations with consistent cancer cell fractions across many samples and determine the cancer cell fraction (CCF) posteriors for each cluster; (ii) *BuildTree* - construction of an ensemble of likely phylogenetic trees and estimation of cancer cell population sizes while accounting for uncertainties in cluster identity and membership; (iii) *GrowthKinetics* - modeling growth dynamics and rates of individual clones while using CCFs, trees and overall tumour burden data; (iv) *CorrectBias* - adjustment of clustered CCF values and cluster sizes across multiple samples in subclones with low CCF to correct for detection biases (correction of “winner’s curse”-like effects); (v) *SinglePatientTiming* - probabilistic reconstruction of patient-specific tumour developmental trajectory, using WES or WGS data from multiple/single samples, including ordering and relative timing of clonal copy-number events, somatic mutations and whole-genome doubling; (iv) *SubclonalTiming* - timing of events within the subclonal branches of the phylogenetic tree; (vi) *LeagueModel* (Cohort Timing) - joint modeling of average progression trajectories from cohorts of patient tumour samples with probabilistic integration of single patient trajectories; (viii) *ConditionalTiming* - identification of statistically significant differences in timing and event trajectories between subsets of samples; (ix) *PhylogicSim ClonalStructure* - simulator of realistic single- and multi-tumour sample data with complex subclonal relationship for evaluating the performance of methods for clustering, tree building, modeling of clonal dynamics and determining fractions of clones in each sample (x) *PhylogicSim Timing* - simulator of cancer progression trajectories and ordering, including realistic single- and multi-tumour sample data, that allows validation of methods for predicting both single-sample and cohort-level trajectories and order of events.

**Figure 1.**
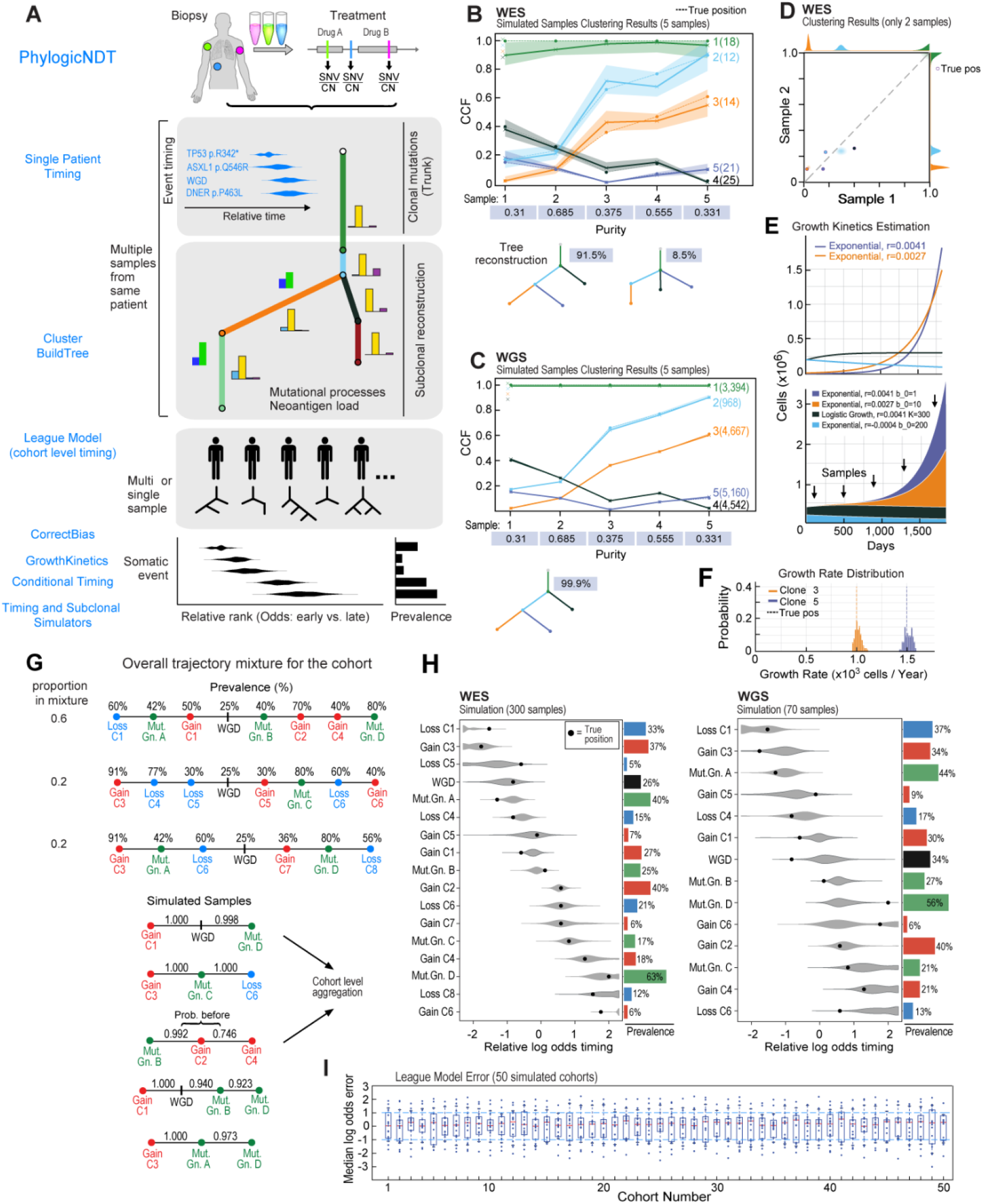
Overview of *PhylogicNDT* analysis of multiple samples from the same patient, and clustering and timing results on simulated whole-exome (WES) and whole-genome (WGS) sequencing data illustrating functionalities of individual tools. (**A**) Schematics of the *PhylogicNDT* suite of tools used to estimate order of mutational events, reconstruction of subclones and their phylogenetic relationships, and comparison of the developmental trajectories and the timing of acquisition of somatic events (early vs. late) across different individuals or subtypes of a disease. Representation of (**B-C**) clustering, (**E-F**) growth kinetics and (**G-I**) timing methods in *PhylogicNDT* for both WES and WGS using simulated data.

### Identification of subclonal composition from multiple tumour samples

Cancer cell populations that develop during tumour progression are inhomogeneously spread across tumour sites and metastatic lesions, while all are genetically related. This creates a variability of abundances of subclones across samples, both in cases when the same lesion is sampled multiple times, or when samples are taken from anatomically distinct lesions ^23^. This intra-sample heterogeneity is even more pronounced in cases when different sampling techniques are used for collecting tumour material, e.g., slices from large tumour resections, core needle biopsy, or sampling of circulating tumour DNA. If sampling occurs along longer time frames - clonal selection, competition, and dynamics also lead to significant differences in abundance of subclones. By jointly analyzing multiple related tumour samples, we can reconstruct their genetic relationships and the evolutionary paths that subclonal populations took to achieve such genetic diversity.

DNA sequencing data (e.g. WES or WGS) allows us to detect allele frequencies of specific somatic events in a sample and, when adjusted by local copy-number and purity, estimate the fraction of tumour cells (CCF) that harbour such a somatic event. Samples can be obtained at different time points or from distinct physiological locations. Cell populations across multiple tumour samples from the same patient (Figure 1A) harbour shared somatic events, and each distinct clone can usually be associated with multiple somatic mutations (both driver and passenger events). Analyzing multiple samples from the same patient provides several opportunities, and hence greater power, to distinguish sublones, (Figures 1B-D) and, if samples are taken at different times, improved ability to monitor their dynamic behavior.

We have developed methods, *PhylogicNDT Clustering* and *BuildTree*, to simultaneously analyze information provided by all the samples from an individual patient. The method utilizes a multi-dimensional Dirichlet process (Figures 1A-C) to jointly estimate the cell population structure and the genetic phylogeny across all samples. The methods are implemented as a two-step process: First, to avoid any bias from artificial or incorrect constraints, we estimate the number of clones and the posterior of their cancer cell abundances and in the second step, we add the tree constraints to estimate the ensemble of most probable phylogenetic trees.

*PhylogicNDT Clustering* takes absolute copy number profiles, purity values and joint mutation calls (**Online Methods**) from all tumour samples belonging to a specific individual. The method can also utilize posteriors distributions on CCF values associated with each mutation across all samples (produced by methods such as ABSOLUTE ^24^). These non-parametric distributions are then subject to a Dirichlet Process (DP) where the distributions over the number of clusters, the CCF value of each cluster in each sample, and assignment of mutations to clusters are sampled via a Markov chain Monte-Carlo (MCMC) Gibbs sampler. Specifically, after initializing a DP with priors as previously suggested ^26,45^, each mutation is first assigned to its own independent cluster. Then, at each iteration of the MCMC, the sampler removes one of the mutations from its cluster and probabilistically assigns it to an existing or a new cluster according to a multinomial probability of joining or creating each cluster. The probabilities are computed across all tumour samples (dimensions) according to the expressions provided below.

The probability of joining each of the k clusters:

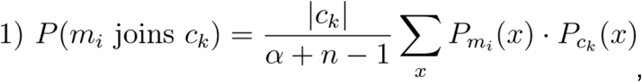

probability of opening a new cluster:

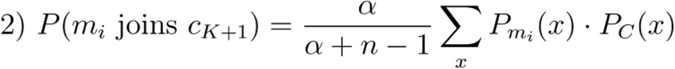

where the posterior of the CCFs associated with each cluster at any given iteration are computed from the multidimensional non-parametric CCF distributions of mutations that belong to the cluster. At the end of a single MCMC round, the *α* parameter, which controls the likelihood of opening a new cluster, is resampled from the posterior given the number of clusters and mutation assignments. The initial priors on *α* used in the method translate into a soft prior on the number of clusters (k), that has a limited impact, if any, on the final result (**Online Methods**).

After completing the MCMC process and discarding the burn-in iterations, we calculate a posterior on both the number of clusters and the N-dimensional CCF distributions of every mutation based on the average of the CCF distributions along the MCMC chain. The multidimensional CCF posterior distributions for each mutation are then hierarchically clustered based on their similarities. If a hard assignment of mutations to clusters is required for downstream analysis, a number of clusters is chosen from the posterior that is consistent with the least complex solution (i.e., fewer clusters), but has at least 10% posterior probability. Finally, the probability that a mutation belongs to a particular cluster is calculated from the CCF posteriors of the cluster and the mutation (**Online Methods**).

### Construction of an ensemble of phylogenetic trees

To generate the ensemble of possible trees that represent the phylogenetic relationships of individual cancer cell populations in the patient, the results and uncertainties generated in the previous *PhylogicNDT Clustering* step are subject to another probabilistic algorithm, called *BuildTree*. The *PhylogicNDT BuildTree* method employs an MCMC Gibbs sampler that assembles likely tree structures by moving individual tree branches (subclones) within each iteration according to a multinomial probability of the tree branch (subclone) being integrated into a specific position within the tree. The multinomial probability is calculated based on the pigeonhole rule (i.e., the sum of CCFs of sibling clones cannot exceed the parent clone’s CCF), while accounting for the uncertainty in assignment of mutations to sub-clones (Figure 1B-C). Specifically, if the new position of the clone implies a parent/child or a sibling relationship, the concordance with such relationship can be estimated through a convolution of the posterior CCF distributions associated with the respective clusters. The convoluted CCF distribution of all children of a specific clone cannot exceed the posterior CCF distribution of the parent clone, and thus we use the probabilities of not exceeding the parental clones in all possible positions to guide the multinomial sampling. This procedure also orders clones in the more likely parent-child relationship based on their CCF distributions (i.e., parent CCF is greater or equal to the child CCF).

Since the CCF posterior distributions from the *PhylogicNDT Clustering* step can partially violate the trees generated in this step (although probabilistically penalized), we also allow mutations to shift cluster assignments after completing the moves of the clusters within the tree. To ensure that the cluster identities and their overall number are stable, we use the CCF posterior from the previous step, as priors in this mutation reassignment step. As a result, the assignment of mutations and the final CCF distributions of each cluster are also influenced by the likely tree structures (**Online Methods**).

At the end of the *PhylogicNDT BuildTree* step the ensemble of probable trees is generated, including the most likely tree, posteriors on the abundances of clones in all samples, and the structure of the cancer cell populations. The multiple samples from the same patient allow for accurate reconstruction from WES or WGS data (Figures 1B-C)

### Probabilistic Estimation of Signature Activities and Neoantigen Load in Individual Subclones

*PhylogicNDT BuildTree* provides an ensemble of probable trees and the uncertainty of the mutational membership in each of the clones in a selected tree (e.g., the most likely one). This probabilistic assignment of mutations to clones can be used to calculate average mutational properties of each clone. One example of such a property is the clone-level activity of mutational processes. These can be used to estimate the differences in the activities of the mutational process across cell populations. We utilize the output of *SignatureAnalyzer*^46^ that annotates each mutation with the probability of belonging to a specific mutational process (based on its mutational signature). We therefore can integrate the uncertainties in the tree and the mutational assignment to tree branches to calculate the posterior activity of the mutational signature for each clone. Similarly, any changes in neoantigen load between clones can also be assessed based on the soft assignment of mutations and thus we can calculate the expected ratio of the likely strong neoantigens to the number of nonsynonymous mutations (Figures 1A and 3; Online Methods). Changes in this ratio may indicate immunoselection of specific clones.

### Modeling of subclonal growth dynamics of tumour cell populations

Accurate estimation of the subclonal structure and phylogenetic tree, accompanied by the uncertainty of such reconstruction, allows us to estimate the subclonal growth dynamics and rates (and their uncertainties) in studies where estimates of overall tumour burden are available in addition to the DNA sequencing data (e.g., white blood cell counts in hematological malignancies, volumetric measurements from radiographic data, and potentially circulating tumour DNA yield from blood biopsies in case of metastatic solid tumors). *PhylogicNDT GrowthKinetics* module integrates information obtained from the *Clustering* and *BuildTree* methods along with the tumour burden measurements to model subclone-specific growth kinetics and estimates growth rates (e.g., in case of exponential growth patterns). The uncertainty in cluster CCF values within the tree is propagated to the estimates of the total number of cells in each of the subclonal populations, which are then used to estimate the possible growth patterns (Figure 1E-F). Comparing growth dynamics of clones to the parental clones enables us to estimate clone-specific fitness.

### Correction of detection bias for low CCF subclones

It is important to appreciate that methods for detecting somatic mutations will not detect a large proportion of low-allele fraction somatic variants. Most algorithms cannot reliably identify mutations with fewer than 3 supporting reads due to intrinsic error rates associated with next-generation sequencing. This inherent limitation in detecting low-allele fraction mutations affects our ability to correctly estimate the posterior CCF values for clusters with mutations close to the detection limit. Thus, often, clusters lying below 15% CCF (depending on purity) in regular WES or WGS data would miss a significant portion of their mutations (those that happen to have lower allele fraction) and, if not accounted for, the posterior on the cluster CCF could be significantly shifted towards higher values. In multi-sample cases, when joint mutation detection is used across all samples, this posterior truncation is not straightforward to estimate given different purities, copy-number profiles, coverages, and histories of the samples. Thus, to account for this specific detection bias across single or multiple samples from the same patient, we have developed a method, *PhylogicNDT CorrectBias*, that utilizes sample-specific coverage and copy-number profiles, mutation data, and clonal structure to estimate the effect of this statistical truncation and adjust the estimates of the CCF values of low-lying clusters towards their true levels. *PhylogicNDT CorrectBias* uses the above sample-specific data to iteratively generate simulated mutations for the cluster, starting with the observed CCF and number of mutations, and gradually lowering the CCF and then introducing the necessary undetected mutations, until the number of detected simulated mutations and their cluster CCF matches the observed values (see Online Methods).

### Simulation of realistic tumour heterogeneity data and validation of clonal reconstruction algorithms

To evaluate the efficiency and accuracy of methods for reconstructing the subclonal architecture and building phylogenetic trees, it is important to have access to high quality truth data. While obtaining extensive experimentally validated data can be challenging, accurate subclonal sample simulations are possible given well-defined principles of how DNA from individual cell populations would be mixed during bulk DNA extraction and sequencing. We have developed a comprehensive method to simulate whole-exome and whole-genome sequencing data on the mutation level that can be used to evaluate methods for subclonal reconstruction, tree building, and estimating subclonal kinetics (Figure 1B-D, Online Methods). *PhylogicSim ClonalStructure* method utilizes purity distributions and coverage profiles obtained from large cohorts of real samples to simulate clonal trees and mutation data for multiple tumour samples from the same patient. Individual samples can have distinct coverage values, purity, and subclonal abundances to represent real-life multi-sample sequencing experiments. This method also allows to encode various growth patterns from individual clones that could then be used to evaluate the performance of tools estimating clonal dynamics and kinetics, such as the *PhylogicNDT GrowthKinetics* method.

### Molecular time ordering of mutational events within a single patient’s cancer

As discussed above, individual patient’s cancer develops on a specific trajectory, i.e. it accumulates somatic events in a defined order that over time leads to malignancy and eventually to broader metastatic spread. If one had an ability to observe the mutations as they occur, thus obtaining a deterministic order and timing of individual events, these trajectories would be well-defined for each cancer cell in a patient, and the trajectory of all clonal events would be shared across all cancer cells. Unfortunately, in the majority of cases, tumours are only detected at a later stage when most of the transforming events have already occurred. By using computational trajectory reconstruction methods that attempt to estimate the most likely order of events from sequencing data, we have a limited probabilistic view into these earlier stages of tumour development. Even though each tumour undoubtedly has its own specific developmental trajectory, it is possible that a large fraction of tumours share a similar path to cancer, reflected in the average or typical trajectory. To explore the trajectories of individual patients and similarities across patient cohorts, we developed a method, *PhylogicNDT SinglePatientTiming*, that uses whole genome or whole exome data from single or multiple biopsies to estimate the order and relative timing of the patient’s somatic events in a probabilistic manner.

As mentioned before, we can use copy number and mutation data together to infer the relative ordering of somatic events in the pre-malignant and early stages of disease in the following ways. When a chromosomal segment is gained, it inherently co-amplifies all somatic mutations acquired within the segment up to that point in time, and thus the relative timing of genomic gains, including whole genome duplication (WGD) events, can be inferred by comparing frequencies of these co-amplified somatic mutations. Using the timing of gains and WGD, small somatic events (e.g., indels and SNVs) and structural variants, can also be timed relative to other events across the genome according to their multiplicity status (the number of chromosome copies that harbor the mutation). WGDs present a unique opportunity to time events across physically disconnected regions of the genome, and provide the ability to compare the timing of mutations, copy number losses, and gains relative to a shared genomic event. Furthermore, subclonal somatic events obviously occur after all clonal events. Indeed, it is possible to infer (with uncertainty) the order of genetic events (i.e., the genetic trajectory) from a normal cell to the sampled tumour state by combining the above information.

The availability of multiple samples from the same patient allows for more accurate timing of events within the patient’s tumour developmental history. For events that are detected across multiple samples, the timing estimate is more accurate due to the multiple measurements of the multiplicity of the detected somatic mutations. For events that are present only in a subset of samples, such information allows to place events accurately in the overall developmental trajectory or, in specific cases, time the occurrence of an event relative to other events within the same subclonal expansion period.

We utilize a relative timing measure, *π*, as defined for every copy-gain in Purdom et al.^31^, which represents the proportion of mutations per unit length of DNA that occurred before the event relative to the total number of mutations on the same DNA interval. Since mutations accumulate over time, this measure can be used to evaluate the “timing” of a mutational event in a hypothetical molecular “clock”. To estimate *π* for any single event, such as a chromosomal gain, we calculate the posterior given a uniform prior on *π*. Effectively, we need to calculate the ratio of mutations that happened before the gain to all mutations that happened before and after the gain (normalization to the amount of DNA at risk for mutations), but also correct the value for any effects associated with the power to detect mutations along the genome and across samples (which depends on purity, coverage and copy-number along the genome). Practically, we can detect individual SNV events along a gained region, and these mutations can either be present after the event, on a single copy of DNA spanning the region (i.e. multiplicity = 1) or after, multiple copies (multiplicity >1). Inherently, the accuracy of timing depends on the amount of data available to estimate these values. Whole-genome data contains significantly more mutations than whole-exome data, thus providing greater accuracy on *π*. Nevertheless, whole-exome data can provide enough mutations to have a wide but appropriate estimation, that can become even more accurate when a single mutation is detected across multiple samples from the same patient. Through integrating posteriors of each mutation, we can calculate and power-correct the posteriors on the proportion of mutations from high and low multiplicity events present on a chromosome segment and thus the *π* of the segment (after change of variables, **Online Methods**).

For higher-order copy number events (i.e., when one or both alleles have a copy number above 2), it is possible to estimate the relative timing of each of the gain events independently via MCMC (see below and **Online Methods**). It is possible that gains that lead to allele copy number >2 happen gradually or in a single burst event. Alternatively, such gains may have occured independently from each other with significant time passing between the first and other subsequent gains.

Overall, *PhylogicNDT SinglePatientTiming* method provides single-patient *π* estimates for all measurable events by directly inferring posterior distributions on all gains, including higher-order gains and WGD, and through convolutions of these posteriors, it also provides *π* estimates for SNVs, indels, copy-neutral loss-of-heterozygosity events, and homozygous deletions.

### Ordering of mutational events within subclones of a patient

In studies of post-treatment tumours, it is quite common that a pre-treatment sample might not be available. In such situation standard genomic analysis that relies on measuring differences between pre- and post-treatment samples is not possible. *PhylogicNDT SubclonalTiming* provides functionality to estimate molecular time ordering within specific branches (including subclonal branches) of the patient’s phylogenetic tree. Given a sufficiently high subclonal mutational burden in a post-treatment tumour (or WGS data), we can determine the order of events and in particular those that happened during the latest subclonal expansions, which more likely have been acquired or selected-for during treatment. In the case when both pre- and post-treatment samples are available, we can more accurately estimate the order of events during progression under treatment and better nominate events that promoted resistance (see below and **Online Methods**).

### Estimating the average cancer developmental trajectory across a cohort of patients

It is widely appreciated that within specific tumour types, individual tumours can be further classified into distinct biological subtypes, based on their histological, genetic, transcriptional, or epigenetic profiling. It could also be hypothesized, based on existing work in premalignant cancer lesions ^5,35,36^, that this is a result of separate developmental trajectories that are specific to the subtype of cancer they define. For example, several major lung adenocarcinoma subtypes were previously identified ^47,48^ and include KRAS-wild type, EGFR-wild type, and fusion-driven tumours. Single patient trajectories can be aggregated together to build preferential timing models, and elucidate the average order of events in a specific subtype. *PhylogicNDT LeagueModel* method ranks the order of driver events based on the order within each patient. The method then calculates the odds ratio of events occurring early (first half) or late (second half) during tumour development (**Online Methods**).

*PhylogicNDT LeagueModel* uses the probabilistic trajectories for single patients generated in the previous step and integrates the information into a pairwise event contingency table (Figures 1G-I). This table represents the probability that a random patient (from the cohort) that harbors a specific pair of events, will have the first event in the pair earlier, later or at a similar/indetermined time as the other event. This dataset is then sampled in such a way that all individual events play a “sports season” against each other with “matches” played between pairs of events, where the outcome of the “match” is decided by sampling from the pairwise probabilities.

To test the ability of the *LeagueModel* to faithfully represent the average trajectory in a cohort we used simulated data. As shown in Figures 1H-I, when a mixture of multiple trajectories is simulated in the cohort (using *PhylogicSim TimingS imulator)*, the *League* model estimates the average trajectory accurately with respect to the relative abundance of trajectories and prevalence of events. To further validate the approach, we simulated 50 distinct cohorts with various combinations of trajectory mixtures, both WES and WGS data. On average, the *PhylogicNDT LeagueModel* accurately estimated cohort-level trajectories (for the majority of the simulated events, the absolute difference between the median of observed and expected log10 odds ratios was below 1, Figure 1I).

### Analysis of differences in developmental trajectories conditional on clinical variables or presence/absence of specific driver events

It is important to understand if known biological subtypes within the tumour show distinct developmental trajectories during progression to malignancy. For this purpose, we have developed an approach, *PhylogicNDT ConditionalTiming*, that allows exploration of differences in progression models between subsets of samples in the cohort. For example, known clinical subtypes of lung adenocarcinomas (EGFR-mut, KRAS-mut, fusion-driven cancer) can be analyzed separately by the *PhylogicNDT LeagueModel* approach and then compared to each other by calculating the difference in the posteriors of their log odds ratio (see below). Significance is calculated using an empirical background model based on permuting samples between the compared subtypes.

Another interesting application of this approach is to explore the association between the absence or presence of a specific mutated gene and the developmental trajectory. Similar to above, a cohort can be split into samples that contain or do not contain a specific event, and any differences in trajectories can be assessed for statistical significance. When performing this analysis across the major driver events in a cancer type, it allows us to identify events that influence the timing of other events (see below and **Online Methods**).

### Exploration of clonal dynamics and progression models of Lung adenocarcinomas before, on and after treatment

To understand the mechanisms and genetics behind tumour progression under multiple lines of treatment and to explore potential trajectories tumours can take prior to diagnosis and during treatment, we applied the *PhylogicNDT* suite of tools to study a subtype of lung adenocarcinomas that are primarily driven by gain-of-function *ALK* fusion events. We then compared the results to a wide range of primary lung adenocarcinomas ^47,48^.

We obtained biopsies from 21 patients, taken at multiple time points and across different anatomical sites, including before, during, and after treatment with 1st and next generation TKIs (total of 64 samples). We sequenced 64 whole-exomes with at least one pre- and one post-treatment samples and matched normal control. In one patient, we additionally had an autopsy analysis at death with total of 13 distinct biopsies analysed.

### Description of the cohort and clinical results

We identified 21 patients with ALK-positive non-small cell lung cancer (NSCLC) who had at least one biopsy available for analysis. Baseline clinical and pathological features of this cohort are summarized in Supplemental Table 1. Median age at diagnosis was 48 years (range 22-77). A majority of patients (85%) had advanced-stage disease at initial diagnosis. Consistent with prior reports ^49^, most ALK-positive patients were never-(80%) or light-smokers (10%), and all had baseline adenocarcinoma histology. Details regarding treatment histories and clinical outcomes are summarized in Figure 2A and Supplemental Table 1. Of note, 81% of ALK-positive patients within this cohort initially received cytotoxic chemotherapy prior to ALK-directed therapies, which largely reflects that most patients (76%) were diagnosed with metastatic NSCLC prior to the first regulatory approval of an ALK inhibitor, crizotinib, in the United States (in 2011).

**Figure 2.**
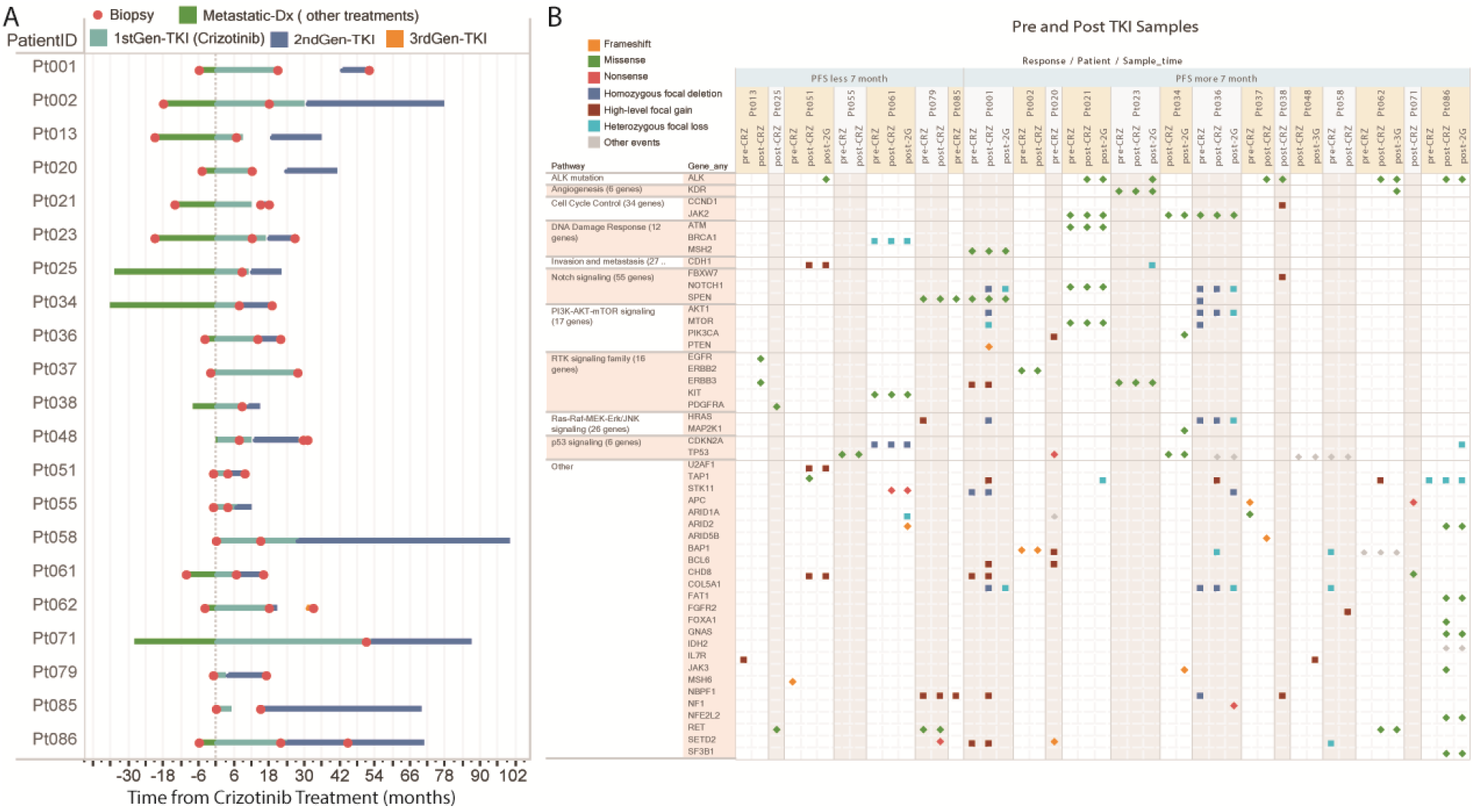
Clinical description and treatment summary of 21 patient ALK-fusion positive non-small cell lung cancer cohort and selected somatic mutations found in their genomes. (**A**) Treatment history plot detailing the clinical treatment history of each patient in the ALK-fusion positive cohort treated with 1st, 2nd, and 3rd generation ALK inhibitors overlaid with time of biopsy (circles). (**b**) Mutation co-occurrence (CoMut) plots showing somatic changes in copy-number and non-silent somatic mutations between pre- and post-treatment samples in known cancer genes and pathways.

All patients (N=21) received crizotinib during their disease course. Median time from metastatic diagnosis to initiation of crizotinib was 5.9 months (range 0.4 – 37.3 months). Patients received a median of one line of therapy (range 0-5) prior to crizotinib. Median progression-free survival (PFS) on crizotinib was 7.9 months (range 2.57-51.6 months; Figure 2A). All patients eventually relapsed on crizotinib. Repeat biopsies performed at or shortly after clinical progression on crizotinib were performed in 19 patients. WES of post-crizotinib biopsies revealed secondary ALK resistance mutations in 5 (26%) patients. The remaining patients did not have ALK resistance mutations, suggesting alternative mechanisms of resistance.

Following disease progression on crizotinib, 20/21 (95%) patients subsequently received a next-generation ALK inhibitor. Agents included: ceritinib (N=14), alectinib (N=3), brigatinib (N=3), and lorlatinib (N=2). Three patients received two or more next-generation ALK inhibitors. Clinical outcomes for each agent are summarized in Figure 2A. In total, 10 biopsies performed on or shortly after progression on next-generation ALK inhibitors were available for WES. ALK resistance mutations were identified in five (50%) of these specimens. Of note, two of these patients (Pt023 and Pt051) were negative for ALK resistance mutations in post-crizotinib/pre-next-generation ALK inhibitor biopsies, consistent with prior reports suggesting that the acquisition of additional ALK resistance mutations is more common in patients treated with more potent next-generation ALK inhibitors.

### Landscape of acquired somatic events after TKI treatment

Upon initial analyses of the 64 samples, we noted considerable changes in copy number variations and acquired mutations of known oncogenic drivers across many well-known cancer gene pathways (Figure 2B). Notably patients that had very short or no response to crizotinib (PFS < 7 months) showed no detectable ALK gene mutation in the post crizotinib biopsy suggesting a certain level of intrinsic resistance, unlike the patients with longer reposes. Individuals that did not acquire known ALK gene alterations post treatment had developed point mutations and copy number events in PI3K-AKT, RAS/RAF/MEK/ERK, RTK signaling pathways. Several patients showed alterations in TAP1 and other genes associated with antigen presentation.

### Tumour progression and clonal dynamics under multiple lines of treatment

A major underlying reason for the emergence of tumour resistance to TKIs after treatment is the acquisition of new, secondary mutations in *ALK*. Accordingly, we closely examined changes in subclonal structure (from *PhylogicNDT Clustering)* that occurred after the TKI treatment(s), specifically searching for growing clones that acquired mutations in *ALK*. We observe that treatment with TKIs is accompanied by strong shifts in the clonal structure (Figure 3). Almost invariably, the resistant clone containing the *ALK* mutation took over and became the majority of the tumour cell population. In all such cases for which the pre- and post-TKI treated samples were available (and had sufficient purity), the post-treatment biopsy contained one or more growing cell populations with an ALK missense mutations.

**Figure 3.**
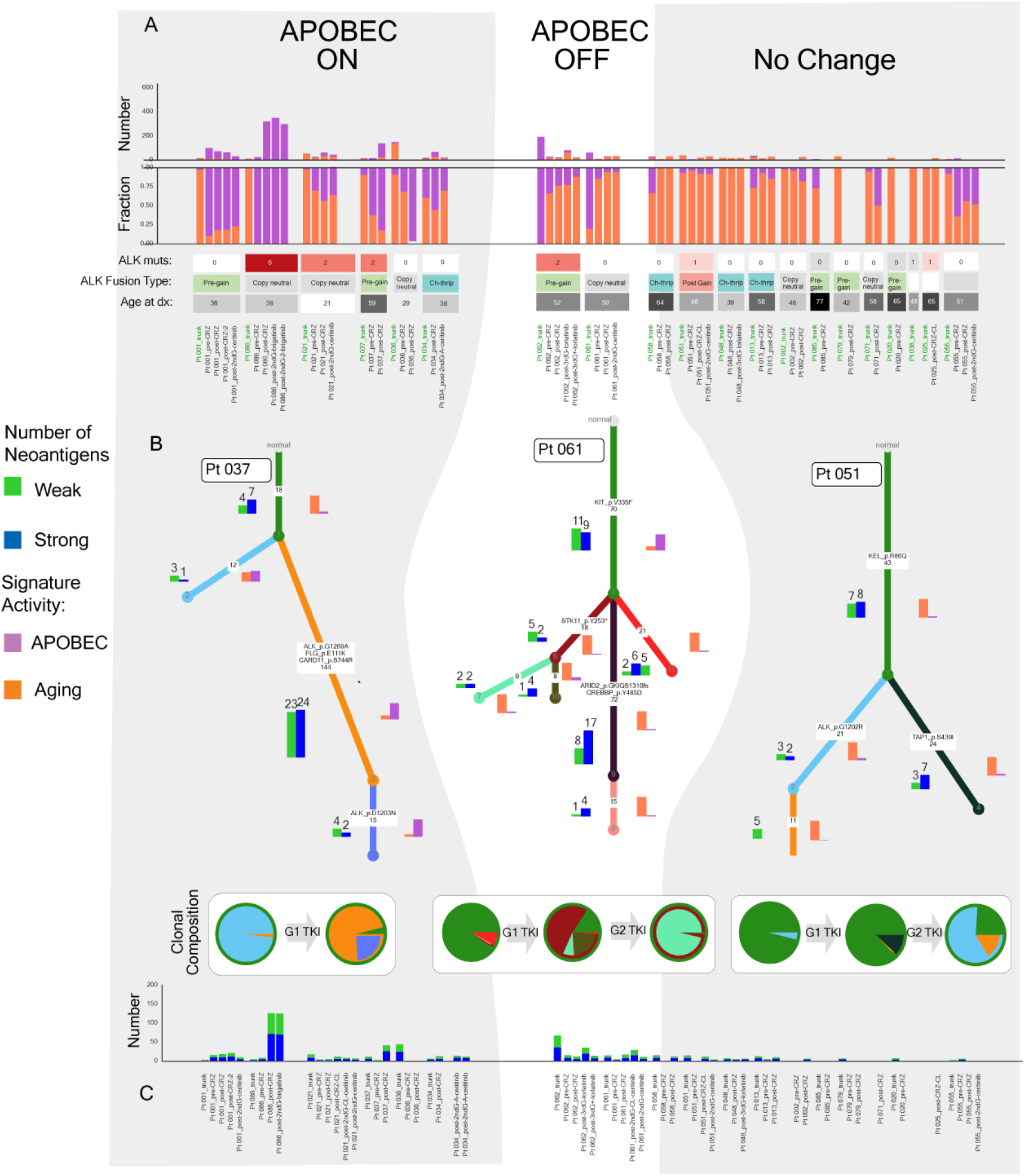
Estimated phylogenetic trees for ALK-fusion positive tumours and subclone identification annotated with somatic mutations, neoantigen number and quality, and predominant mutational signatures in each sample. (**A**) Signature analysis of mutational signatures active in each sample by the *SignatureAnalyzer* tool. (**B**) Phylogenetic trees built by *PhylogicNDT BuildTree* representing clonal structure of each tumour after integrating data from multiple samples from a single patient, overlaid with signature analysis and neoantigen load for each clone and subclone. Inferred clonal composition of the pre- and post-treatment samples is represented as circle plots. (**C**) Neoantigen number and fraction of weak (green) vs. strong (blue) antigens in each sample.

Mutations can be usually attributed with some probability to a particular mutational process active in a cell (e.g., normal aging or deficiencies in particular repair processes). To determine which processes might have been responsible for mutating *ALK* under pressure of TKI treatment, we used our *SignatureAnalyzer* ^46^ tool to discover the signatures and their level of activity in each sample from each patient and annotated each *ALK* mutation with the probability that it originated from a particular mutational process (Figure 3A-B). Interestingly, the APOBEC (COSMIC SBS 2, 13 ^50^) and aging signatures (COSMIC SBS1) were the most prominent mutational processes found in our cohort.

We constructed phylogenetic trees for each individual patient using the *BuildTree* tool (representative trees shown in Figure 3B) and assigned mutations to each of the branches. We quantified the relative contribution of signatures that were originally active in a patient at the start of the disease, as determined by their common appearance in every sample (i.e. truncal mutations), with those that were active in later cell subpopulations (Figure 3A). In 9 of the 21 cases, we observed activity of a mutational process associated with APOBEC deaminases (>20% APOBEC-associated mutations across all samples of the patient). Moreover, in 6 of these 9 cases, there was a significant increase in APOBEC activity in later cell subpopulations compared to the truncal clone (increased by more than 2-fold; see example in patient Pt 037 tree in Fig 3B). In another 2 cases, the truncal mutations already had APOBEC-associated mutations but later subclones predominantly acquired aging signature mutations, and nearly no APOBEC (e.g. Pt 061 tree); a third category of 12 patients displayed no change in the mutational signatures between the trunk and subclones (e.g. Pt 051 tree).

Next, we tested whether patient-specific factors are associated with signature activities. Most notably, we identified an unexpected significant association between the APOBEC signature activity and patient age. The majority of cases that showed a significant increase in APOBEC activity over time were the relatively young (< 40 yo) patients: nearly all young patients displayed higher APOBEC activity and mutation load (83%, N=5/6) than the older (>= 40 yo) patients (20%, n=2/10) (Fisher’s Exact Test p=0.0350).

As expected, the number of truncal mutations associated with the <u>C</u>pG>T Aging signature (COSMIC SBS 1) in younger patients was consistent with their age (noticeably lower than in the patients above 40 yo). One can hypothesize that the substantially increased activity of the APOBEC signature in younger patients is required for their cancer initiation and progression given the low mutation counts associated with aging. This high APOBEC activity may also influence the patient’s response to TKI inhibition and the tumour’s ability to develop resistance due to the higher genetic diversity which contributes both to intrinsic and acquired resistance. This is supported by a 38-year-old patient (Pt 086) who developed 6 individual *ALK* mutations and had one of the highest APOBEC mutation loads in the entire cohort. In contrast to the relatively younger patients, most older patients displayed a relative absence of APOBEC activation in either pre- or post-treatment samples, and their overall mutation load was appropriate for their age (since most were from the CpG>T Aging signature). Additionally, two patients that exhibited moderate APOBEC activity in the clonal trunk surprisingly showed decreased activity in the later subclones.

Given the substantial differences of APOBEC activity with age in our cohort, we next explored whether other genomic features correlated with age. We found a trend suggesting that the structure and type of ALK fusion rearrangement also differs between younger and older patients. We classified the local copy number and pattern of the EML4-ALK rearrangement into the following 3 categories: (i) copy-neutral inversion, (ii) high level gain of the fused allele and (iii) chromothripsis. Younger patients exhibited more copy-neutral EML4-ALK fusion fusions without associated gains (3/5), while older patients exhibited high chromothripsis events associated with amplification and multiple gains.

### Clonal composition changes during treatment of individual patients

To study changes in clonal composition due to TKI treatment, we analyzed in each patient the sizes of the clones, as identified in the phylogenetic tree, and in each of the samples (circle plots for representative patients in Figure 3B). We noted significant shifts (>%25 CCF shift in at least on clone) that occurred in the clonal composition after TKI treatment in almost every case. Individuals that acquired on-target ALK resistance mutations in one or more subclones showed significant clonal shifts in their post-treatment biopsies, wherein the ALK-resistant cell population overtook a significant portion of the tumour (e.g. Pt 037 and Pt 051 in Figure 3B). In patients where the resistance to crizotinib could not be explained by ALK mutations, we were still able to detect very strong clonal shifts between pre- and post-treatment biopsies, suggesting that the emerging clones harbor alternative resistance mechanisms. Some growing clones acquired mutations in known cancer driver genes (as in Pt 61, with STK11 mutation), but whether these driver mutations conferred resistance or independently increased fitness is yet unclear. In patients that developed kinase-domain mutations after crizotinib, secondary resistance often also was caused by additional events in the ALK gene.

An important clinical question is whether the mutation load increases in samples after several lines of treatment compared to the primary tumour. If this is indeed the case, patients with significant mutation load in their tumours might be considered for treatment with checkpoint inhibitors, since recent results show better response to immunotherapy in high mutation load tumours ^51^. Neoantigens arise from mutated proteins due to nonsynonymous mutations acquired during tumour growth, and their quantity and presentation quality (weak or strong) affect the ability of the immune system to recognize and kill tumour cells ^52^. The number of neoantigens introduced with each clonal expansion during tumour evolution is usually proportional to number of mutations acquired during the expansion ^51,52^. Fewer than expected neoantigens can be a hallmark of resistant subclones that survived the selective pressure of the immune system (ie. immunoediting).

The clonal tree generated by *BuildTree* allows us to assess and quantify the neoantigen load on a clone-by-clone basis (Figure 3B-C). As expected, the neoantigen load was proportional to the number of mutations within the respective clone in most scenarios; thus, clones displaying increased APOBEC mutational activity also had proportionally higher neoantigen load. In some cases, we identified unexpected discrepancies between neoantigen loads in patient’s clones with a similar number of mutations. For example, a clone harboring a likely deactivating mutation in TAP1—a known gene involved in transporting antigens from the cytoplasm to the endoplasmic reticulum—in patient Pt 051 showed a markedly higher number of strong neoantigens compared to its sibling clone (3 vs 12 strong neoantigens among the respective 21 vs. 24 clone-specific mutations; Fig 3B; Fisher’s Exact p-value=0.0071). This behavior can suggest immunoediting at the subclonal population level and can also explain increased fitness of the TAP1-mutated clone in the intermediate post-crizotinib sample. After treatment with 2nd generation TKI, however, the ALK-mutated clone was preferentially selected (Figure 3B).

Overall, by representing the clonal structure with a phylogenetic tree and studying changes in per-sample clonal composition, we were able to explore the differential behavior of driver mutations, mutational signatures, copy number changes, fusion events, and neoantigen load on a clone-by-clone basis and thus reconstruct clonal dynamics under treatment with increasing levels of detail.

### Single-patient timing: Molecular time ordering of mutational events and developmental trajectory from individual patients

We applied *PhylogicNDT SinglePatientTiming* to a large set of WES and WGS data from 455 lung adenocarcinoma patients (union of TCGA data and our cohort) to infer patient-level preferential ordering trajectories. Depending on the tumours architecture and mutational rate, these trajectories can be inferred with varying levels of resolution. We selected a representative set of patients (Figure 4A) from 3 major known clinical subtypes (fusion-driven, KRAS-mut, EGFR-mut) of lung adenocarcinoma to demonstrate typical differences between individual trajectories. As expected, tumours from the EGFR-mut and KRAS-mut subtypes show early EGFR and KRAS point mutations, respectively, followed by the gains in the general chromosomal regions of those genes. Surprisingly, the trajectory of some *ALK/ROS1* fusion-driven tumours showed early *U2AF1* mutations, a gene with a known role in splicing of RNA transcripts, suggesting a possible contribution in promoting growth of cells driven by fusion events.

**Figure 4.**
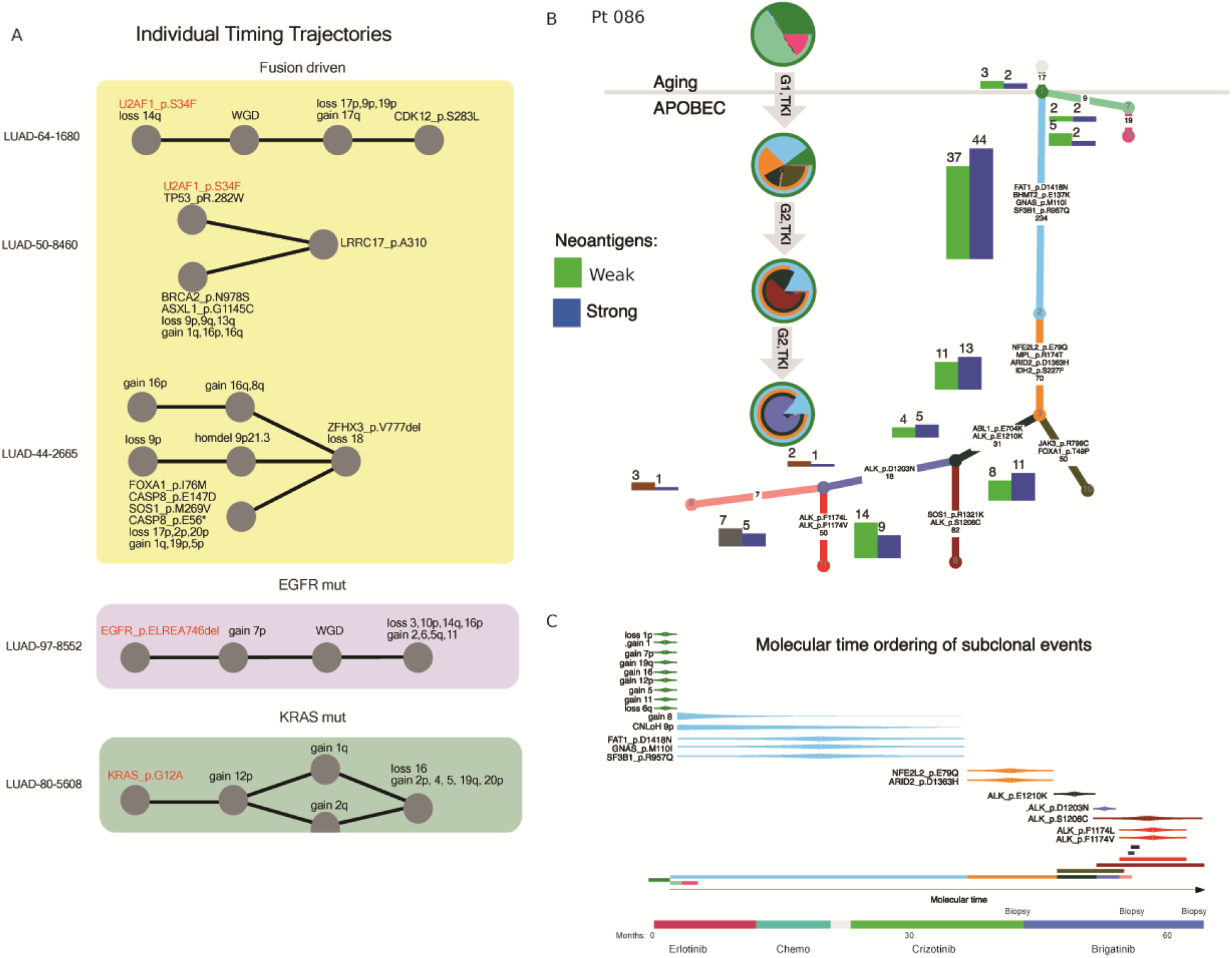
Single-patient timing trajectory: Molecular time ordering of mutational events and timing trajectory from individual patients. (**A**) Individual timing trajectories resulting from *PhylogicNDT SinglePatientTiming* analysis of selected ALK-fusion positive as well as EFGR-mutated and KRAS-mutated lung adenocarcinoma patients. (**B**) Phylogenetic tree and molecular time ordering (**C**) of subclonal events (within a tree branch) by *SubclonalTiming* analysis from a single patient. For molecular time ordering, treatment schedule is shown below to visualize the changes in subclonal mutational events and the acquisition of potential resistance mutations in the context of when the patient received each of the treatments, 1st generation TKI (crizotinib), 2nd generation TKI (brigatinib).

The availability of multiple samples from the same patient enabled both improved accuracy in reconstruction of the single-patient phylogenetic tree and timing of events (see **Online Methods**, Supp. Figure 3). Similar to timing events within the trunk of a phylogenetic tree, it is also possible to time events within a specific branch with the *PhylogicNDT SubclonalTiming* tool, as mentioned above. When events are present only in a subset of samples from the same patient, this information allows to more accurately place events in the overall developmental trajectory within a specified subclonal branch. Moreover, we can estimate the timing of these events with respect to mutations specific to their branch. For example, in Figure 4B, we find that a prominent subclone in Pt086 acquired gains on chromosomes 8 and 9p, and using the other mutations in this branch, we can infer their timing. We observe that prior to TKI treatment, the patient’s mutation rate is dominated by the aging signature (COSMIC SBS 1). Following crizotinib, the mutation rate increased sharply, with a strong contribution of the APOBEC signature, likely driving resistance (consistent with the observation described above), and the neoantigen load increased more than tenfold (2 vs. 44). Using the multiple samples available for this patient, we are able to order the subclonally acquired copy number gains and mutations, showing that chromosome 8 and 9p are potential founding events of this subclone. Additionally, the phylogenetic tree provides information on the molecular time (measured in number of acquired mutations) that different subclones have emerged. For example, since the orange subclones is a descendant of the cyan subclone, we can infer that it emerged after all the mutations in the cyan subclone. The molecular time of sibling subclones start at the end of their parental clone (Figure 4B). We see a multitude of ALK mutation clones occuring in serial and sibling clones in the complex of the phylogenetic structure of this case.

To validate that our timing estimates within a subclone are correct, we reran the analysis after excluding the earliest sample and, in this scenario, the mutations present in the cyan clone were detected as fully clonal (truncal). Reassuringly, the relative timing of the events when the cyan clone is a subclone (i.e., when using all samples) are consistent with the ones obtained when the cyan clone is part of the trunk (i.e., when ignoring the earliest sample) (Supp. Figure 4). This finding is quite important since in many treatment scenarios where only post-treatment samples are available, we could infer the timing and order of events that happened late in development, hence likely occurred during treatment. Additionally, such analysis allows us to explore the molecular time in which neoantigens are introduced relative to different treatments.

### Cohort-level timing analysis: Comparison of developmental trajectories of fusion-driven, EGFR-mut, and KRAS-mut lung adenocarcinoma subtypes

It is widely appreciated that specific tumour types can be further categorized into biologically meaningful subtypes based on their histological, genetic, transcriptional, or epigenetic profiling–for lung adenocarcinoma, these major subtypes include KRAS-mut, EGFR-mut, and *ALK/ROS1* fusion-driven tumours ^53^. It can be hypothesized that developmental trajectories (order and type of events) will differ among subtypes.

To explore and compare the differences in the average evolutionary trajectories that transform normal cells in the specific cancer subtypes, we ran the *PhylogicNDT SinglePatientTiming* and *LeagueModel* tools on individual tumours and known clinically distinct subtypes from a dataset of 455 adenocarcinomas (union of TCGA samples and our cohort).

In Figure 5A, we present the *LeagueModel* timing diagrams for the different subtypes of lung adenocarcinoma. We find that all three subtypes have relatively early *TP53* mutations and chromosomal loss of 17p, though non-fusion driven cancers tend to have them in a set of earliest events. This suggests a shared biological dependency on initially mutating the *TP53* locus for tumour development regardless of the genetic subtype. EGFR-mutant and KRAS-mutant lung cancers show very early co-amplifications of the corresponding gene loci that occur after the initial *EGFR/KRAS* mutation, suggesting a strong tendency during early stages of tumours to amplify and up-regulate the mutated copy of the gene. Figure 5B displays major differences between fusion-driven and all other lung adenocarcinomas that we analyzed. In Figure 5C, we quantified the differences between the three major subtypes of lung adenocarcinoma. The figure depicts three classes of events: events that are 1) significantly earlier or 2) significantly later in fusion-driven cancers compared to either EGFR-mutant or KRAS-mutant cancers (permutation-based; left/right in Figure 5C; **Online Methods**), and 3) events in which the timing is not significantly different in fusion-driven cancers (middle in Figure 5C). As expected KRAS and EGFR driven cancers showed these defining events as substantially earlier in the timelines as well as relatively early gains of corresponding chromosome regions.

**Figure 5.**
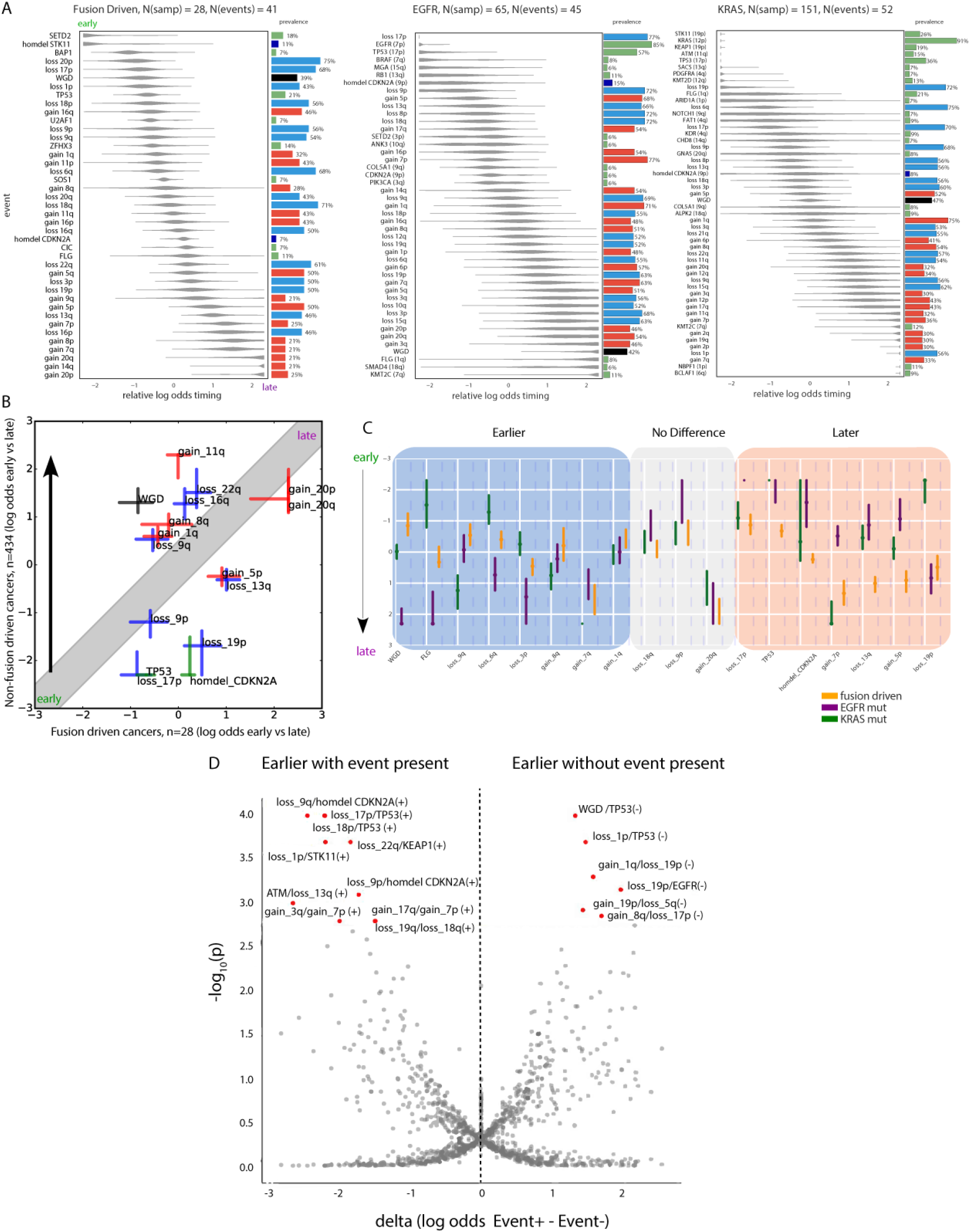
Cohort-level timing analysis: Comparison among the acquisition order of somatic mutations in fusion-driven, EGFR-mutated, and KRAS-mutated lung adenocarcinoma subtypes. (**A**) *LeagueModel* timing diagrams of combined trajectories from a cumulative dataset of 455 patients containing patients from each lung adenocarcinoma subtype showing when in mutational time (early → late) somatic mutations are acquired. (**B**) Comparing the timing of selected acquired somatic mutations between non—fusion-driven and fusion-driven lung adenocarcinoma cohorts, with differential events shown. (C) Quantification of the differences among the three major subtypes of lung adenocarcinoma, divided into three classes of mutational events: events that are 1) significantly earlier (left) or 2) later (right) in fusion-driven cancers compared to either EGFR-mutated or KRAS-mutated cancers and 3) events in which the timing is not significantly different among the three subtypes (middle). The significance values are based on permutation test with the 3 categories. (**D**) “Butterfly” plot of *PhylogicNDT ConditionalTiming* results capturing the association between the presence or absence of specific somatic events and the developmental trajectory, to determine potential biological dependencies of specific late-occurring mutations on other, earlier-occurring events.

Further exploration of the trajectories is possible by calculating conditional dependency of timing of certain genetic events. We explored whether the presence or absence of a specific driver event (mutation or copy-number) significantly impacted the relative timing of any other event. We developed a tool *PhylogicNDT ConditionalTiming* and a permutation test to explore the conditional timing of somatic events (Figure 5D, “butterfly” plot). Known biological dependencies are clearly observed, such as chromosomal loss of 17p locus occurring earlier in TP53-mutated cancers (n=213/236 TP53-mutated, n=104/208 TP53-wild type) and loss of 9p occurring earlier in samples with homozygous deletion of *CDKN2A* (n= 47/49 homozygous deletion, n= 237/395 no homozygous deletion), ATM/loss_13q have similar dependency. In addition, we identified yet-uncharacterized conditional timings that can provide novel biological insights into cancer development, such as earlier *BRAF* mutations in cases with loss of 15q arm (n=22/249 with loss of 15q, n=16/195 without loss of 15q). Finally, loss of 19p appears earlier in EGFR-wild type cases (n=39/59 EGFR-mutated, n=275/385 EGFR-wild type), suggesting that tumours that do not have *EGFR* alterations are likely driven by loss of *STK11*.

### Heterogeneity of tumour lesions in anatomically distinct locations

We applied the whole *PhylogicNDT* package to multiple samples taken at the same time from various metastases across the body (lung and liver) from the same patient at autopsy. By applying *PhylogicNDT* to this autopsy case, we were able to assess the phylogenetic relationships and migration patterns of clones in a very large sample set from a single patient. We analyzed 13 samples from patient Pt 034 (Figure 6). The spatial heterogeneity of the disease is quite remarkable when the clonal structure on individual samples is mapped back to their location in the patient’s body (Figure 6). We observe that the lung and liver lesions are markedly different: with one branch (which first showed progression) driven by a known resistant *MAP2K1* missense mutation and the other likely driven by *JAK2* and *PIK3CA* missense mutations. The *MAP2K1* branch is associated with the Aging mutational signature (the most active signature in all lesions), whereas the *JAK2* branch with lesions in the liver show high levels of APOBEC activity. The two branches have distinct copy number events: the first branch harbors gains in 14q and 3q, as well as loss of 13q, while the second branch is associated with loss of 1q.

**Figure 6.**
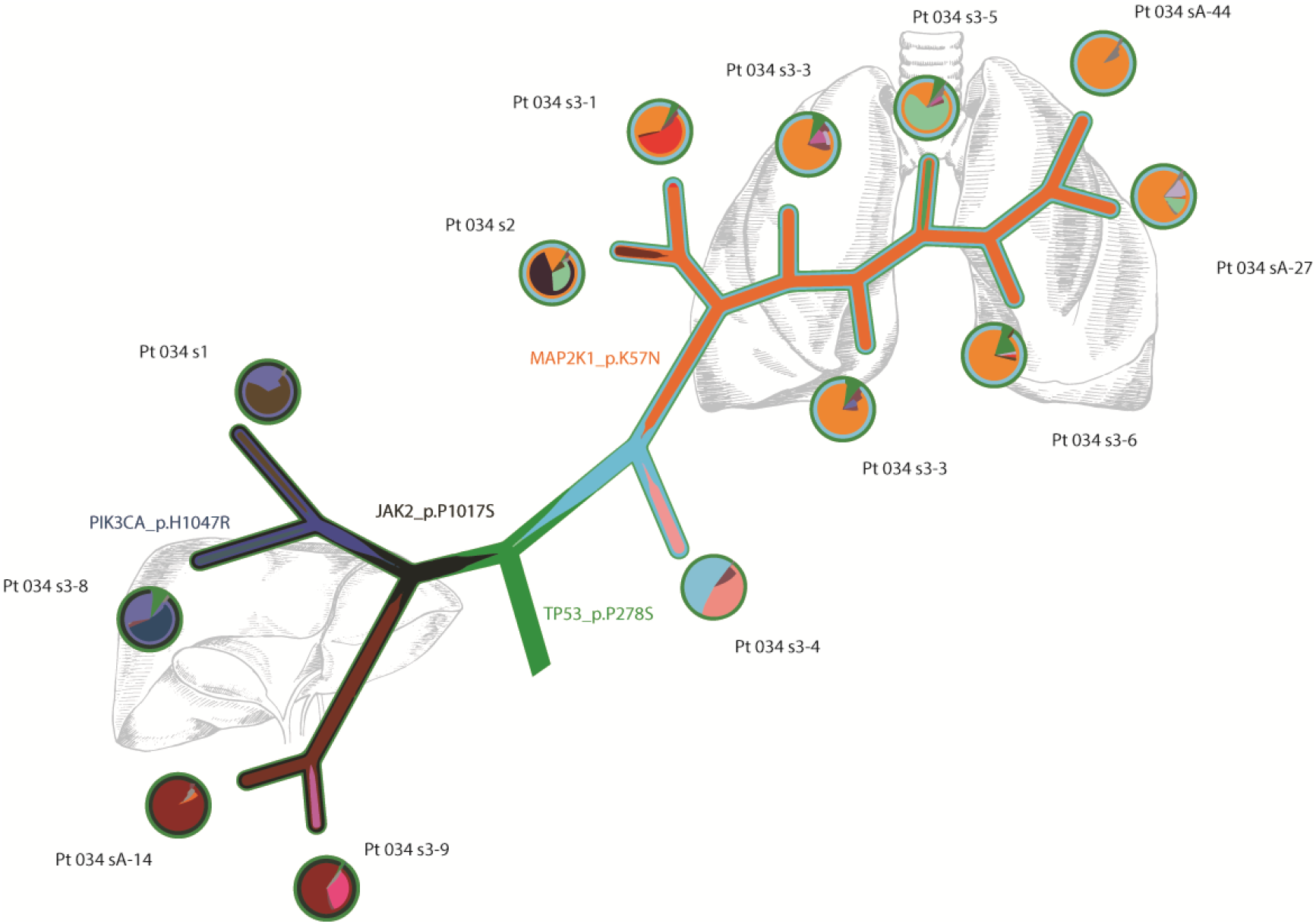
Heterogeneity of tumour lesions in anatomically distinct locations, sampled during treatment and time of autopsy of an ALK-fusion positive lung adenocarcinoma patient (Pt 034). Clonal migration (“River”) plot detailing subclones migrations between 13 samples. Truncal clone is depicted in green, with branches representing major subclones. Left side are subclones found in 4 liver lesions. Right side of truncal clone are subclones found in 9 lung lesions. Circles represent the fraction of each subclone population found in each sample. Known ALK inhibitor resistance mutation in MAP2K1 found in lung lesions (orange clone).

This finding highlights the anatomical aspect of intratumour heterogeneity, showing that different anatomical sites (liver vs. lung) can harbour lesions with different activity of mutational processes (which may be related to different microenvironments) that result in a different probability of developing resistance. Interestingly, the lung lesions also show evidence of cross-lesion seeding, since multiple subclonal events are shared among lesions across both the left and right lobes of the lung.

## Discussion

The development of tumour resistance to therapeutic interventions remains a key roadblock to complete and sustained tumour eradication from the host. Thus, understanding the history of tumour progression and growth at all stages of disease (premalignant, primary, post-therapy, and upon recurrence) is key to understanding resistance. Here, we presented an analytical suite of tools, *PhylogicNDT*, capable of statistically reconstructing the history of tumour growth and progression by analyzing multiple samples from the same patient. To demonstrate the application of *PhylogicNDT*, first and next generation TKI-treated ALK lung adenocarcinoma samples from multiple timepoints were analysed to better understand tumour dynamics before and after treatment. We were able to explore dynamic changes of tumour cell populations as well as interpret the order of somatic events as they occur during tumour initiation and progression. We compared the progression models of distinct molecular subtypes of lung adenocarcinomas and found both expected and novel order dependencies in these cancers.

Several groups, including our own, have previously developed mathematical models and computational approaches to explore clonal structure of cancer cell populations and compare samples from an individual patient to each other ^5,6,8,9,19^. Additionally, approaches were developed to analyze mutational multiplicity rates mostly from primary tumours in an attempt to order clonal events that preceded the last detectable clonal expansion (including early mutations that possibly occurred before malignancy). We significantly expanded our earlier approaches ^5,7,9,19,26^ to be able to jointly model both clonal dynamics and order of early events in cancer across a large number of samples representing the same patient. This allows us to explore developmental trajectories of tumours not just by using whole genome sequencing and data on single samples, but by integrating multiple whole exome sequences and often at even greater resolution. We also developed a sampling methodology *(PhylogicNDT LeagueModel*) that allows to infer average developmental trajectories by integrating larger cohorts of tumours. This analysis provides more robust results as the cohort size increases and therefore benefit from integrating with both WGS and WES data, generated by large cancer genome projects such as The Cancer Genome Atlas and the International Cancer Genome Consortium, as well as other projects.

We demonstrated the utility and accuracy of the PhylogicNDT suite of tools by evaluating their performance on extensive simulated data and by applying them to 434 lung adenocarcinomas as well as 21 patients with multiple samples (up to 13 samples per patient) collected in different timepoints after treatment. We identified distinct developmental trajectories between clinically known lung adenocarcinoma subtypes -- *KRAS*-mutated, *EGFR*-mutated, and fusion-driven lung adenocarcinomas. For example, *KRAS-* and *EGFR*-driven cancers had not just point mutations in the corresponding genes but also early co-amplifications of the mutated alleles. On the other hand, less common drivers tended to occur preferentially early in specific tumour subtypes. When our results are compared to previously reported models of progression from pre-malignant to malignant lung cancer (based on sampling pre-malignant lesions) ^54^, we find similar patterns.

Our results from the joint analysis of multiple samples from the same patient support previous findings about widespread intra-tumour heterogeneity^23,37,38^, but also suggest incredible plasticity of the tumour cell populations under treatment with strong propensity to select specific clones after recurrence, even when the genetic cause of the resistance cannot be unambiguously determined. By using multiple samples, we were able to reconstruct the phylogeny of all detectable cancer cell populations and characterize the subclones in terms of activity of mutational processes and neoantigens load. Our single-patient timing approaches allow us to explore the order of events not just in the trunk of the phylogenetic tree but also within specific clonal expansions, opening possibilities to differentiate between late, post-treatment events and earlier pre-treatment mutations. Thus, this approach can be used to nominate resistance mechanisms by only analyzing sets of post-treatment samples (such as post-treatment blood biopsies) without the necessity of obtaining primary or pre-treatment tissue (which is often logistically challenging). These approaches, when applied to our cohort of ALK-rearranged lung adenocarcinoma patients, discovered several interesting patterns of mutational signature activity in later subclonal populations. Specifically, we found statistically significant differences in APOBEC activity in younger vs. older patients, suggesting a role of this mutational process in tumour initiation in younger patients. We also demonstrated activity of different signatures depending on the physical location of the metastatic lesions suggest both genetic and, potentially, environmental modulators of mutational process activity in drug-resistant tumours.

The methodology developed in this study can be applied to a broader set of cancers and datasets, including multiple sampling in autopsy cases, time course blood biopsy measurements, and pre- and post-treatment analysis of resistant tumours. Studying both the biology of early tumour development and its progression after treatment is clearly needed to answer the questions of resistance and to find new vulnerabilities and therapeutics for effective treatment of cancer.

## Acknowledgments

G.G. and I.L. are partially funded by the Broad/IBM Cancer Resistance Research Project and NCI (R01DE022087, to J.W.R). G.G. is partially funded by the Paul C. Zamecnick, MD, Chair in Oncology at MGH and NCI CLL grant (P01CA206978-01). This work was supported in part by a grant from the National Cancer Institute (5R01CA164273, to A.T.S.) and an ALK-Positive Lung Cancer Transformational Research Award/Lungevity Foundation (to J.F.G). C.J.W. acknowledges support from NHLBI (1RO1HL103532-01) and NCI (1R01CA155010-01A1, U10CA180861-01), and is a Scholar of the Leukemia and Lymphoma Society. Authors are grateful for useful comments and editing help from M. Miller.

## Software availability

The PhylogicNDT package will be available for download from https://github.com/broadinstitute/PhylogicNDT.

